# White matter microstructure in face and body networks predicts facial expression and body posture perception across development

**DOI:** 10.1101/2022.06.16.494491

**Authors:** Isobel L. Ward, Erika P. Raven, Stephan de la Rosa, Derek K. Jones, Christoph Teufel, Elisabeth von dem Hagen

## Abstract

Facial expression and body posture recognition have protracted developmental trajectories. Interactions between face and body perception, such as the influence of body posture on facial expression perception, also change with development. While the brain regions underpinning face and body processing are well-defined, little is known about how white-matter tracts linking these regions relate to perceptual development. Here, we obtained complementary diffusion magnetic resonance imaging (MRI) measures (fractional anisotropy FA, spherical mean 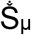), and a quantitative MRI myelin-proxy measure (R1), within white-matter tracts of face- and body-selective networks in children and adolescents and related these to perceptual development. In tracts linking occipital and fusiform face areas, facial expression perception was predicted by age-related microstructural development, as measured by 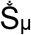 and R1, as well as age-independent individual differences in microstructure, as measured by FA. Tract microstructure linking the body region in posterior superior temporal sulcus with anterior temporal lobe (ATL) was related to the influence of body on facial expression perception, supporting ATL as a site of face and body network convergence. Overall, our results highlight age-dependent and age-independent constraints that white-matter microstructure poses on perceptual abilities during development and the importance of complementary microstructural measures in linking brain structure and behaviour.

## Introduction

The ability to recognize other people’s facial expressions, a critical skill for everyday social interactions, has a protracted development that extends well into adolescence (Calder, 2011; Gao & Maurer, 2009). While functional and structural changes (Gomez et al., 2017; Natu et al., 2016) in face-selective brain regions have been linked to developmental improvements in the perception of facial characteristics such as face identity, our understanding of brain development relating to facial expression perception is limited at best. Expression recognition is subserved by a network of regions in lateral occipitotemporal and ventral temporal cortex (Duchaine & Yovel, 2015), but the developmental changes in white-matter connectivity between these regions and their relation to perceptual development are unknown. Moreover, facial expression perception has to be considered within the context of other cues that are used to recognise emotions. Most prominently, another person’s body posture provides a context that exerts a powerful influence on how facial expressions are perceived (Aviezer et al., 2008; Meeren et al., 2005; Teufel et al., 2019), suggesting possible interaction between face and body networks. In the current study, we therefore investigated the development of perception of facial expressions, body postures, and the influence of body context on expression perception through childhood and adolescence. Additionally, we investigated microstructural changes within white matter connections along face and body processing pathways in the brain that directly underpin perceptual development.

Between infancy and adolescence, humans become increasingly more accurate at perceiving the subtle changes in facial expressions that are associated with different emotions (Herba et al., 2006; Thomas et al., 2007). The development of emotion recognition from body posture has been studied far less, but suggests a similarly protracted developmental trajectory to facial expression perception (Heck et al., 2018; Ross et al., 2012; Vieillard & Guidetti, 2009). At the cortical level, faces and bodies are thought to be processed along largely distinct cortical pathways in lateral occipitotemporal and ventral temporal cortex (VTC). ‘Core’ face-selective regions have been shown to include the occipital face area (OFA), fusiform face area (FFA), posterior superior temporal sulcus (pSTS-Face) and anterior temporal lobe (ATL) (Duchaine & Yovel, 2015; Haxby et al., 2000). Similarly, and in close proximity to face-specific regions, a set of body-selective regions have also been described, including extrastriate body area (EBA), fusiform body area (FBA), and pSTS (pSTS-Body) (de Gelder, 2006; Peelen & Downing, 2005).

A limited number of studies have shown that the selectivity, sensitivity, and/or size of face-selective regions increase during development and are related to facial identity perception (Golarai et al., 2010; Gomez et al., 2017; Natu et al., 2019). For instance, the activity of face-selective regions as measured by fMRI was found to be more strongly modulated by subtle differences in face identity in adults compared to 5-12 year-old children (Natu et al., 2016) and these neural sensitivity differences were related to perceptual performance in discriminating facial identities, even when controlling for age. These findings are consistent with the idea that within-region developmental changes such as a sharpening of tuning functions of face-selective populations of neurons might be directly related to perceptual improvements across development (Natu et al., 2016). While it is currently unknown whether similar developmental changes within face-selective areas might underpin improved perceptual ability in relation to other face characteristics, including facial expression, it seems plausible to assume that this is the case.

A complementary process that likely shapes the development of facial identity and facial expression perception are changes in structural connectivity between cortical regions. The core face network is underpinned by direct white matter connections between face-selective regions along the occipital and ventral temporal cortex (Gschwind et al., 2012; Pyles et al., 2013; Wang et al., 2020). In adults, there is some evidence linking the structural properties of white matter at the grey-white matter boundary close to fusiform-face regions to identity recognition ability from faces (Gomez et al., 2015; Y. Song et al., 2015). However, the extent to which the microstructure of tracts between face-selective regions relate to perceptual ability, in particular facial expression perception, and how the developing microstructure constrains this ability has not been addressed to date.

Research in adults has demonstrated that the body context within which a face is perceived can have dramatic effects on facial expression perception (Aviezer et al., 2008, 2012; Hassin et al., 2013; Meeren et al., 2005; Teufel et al., 2019), suggesting some interplay between face-and body-selective pathways. For example, observers are more likely to perceive a disgusted face as angry when presented in an angry body context. Developmental research suggests that children’s facial expression perception is even more strongly influenced by body context compared to adults (Leitzke & Pollak, 2016; Mondloch, 2012; Mondloch et al., 2013; Nelson & Mondloch, 2017; Rajhans et al., 2016). A changing structural connectivity profile between face-and body-selective areas is one potential substrate underpinning these developmental changes in contextual influence on facial expression perception. Work in both adult humans and primates (fMRI and electrophysiology) suggests that non-emotional face and body information is processed largely independently in the early visual system, with influences of the body network on face processing emerging later in the face-selective hierarchy in ATL (Fisher & Freiwald, 2015; Harry et al., 2016). A recent psychophysical study points to the existence of a similar processing hierarchy for the integration of emotion signals from face and body (Teufel et al., 2019). Microstructural properties of connections from face- and body-selective areas to ATL are therefore of particular interest to understand the development of facial expression perception in the context of an expressive body.

In the current study, using the precision afforded by psychophysical measures, we explored the developmental improvements of facial expression and body posture discrimination in 8–18-year-olds, and their relation to the development of contextual influences of body posture on facial expression perception. Using functionally-localised face-and body-selective seed regions for fibre-tracking in the same individuals, we isolated white matter connections within the face and body networks and their convergence onto candidate regions integrating face and body information, notably ATL. We extracted multiple microstructural measures from these tracts to specifically address the component processes of microstructural change across development and their relation to perception. In particular, we obtained a diffusion magnetic resonance imaging (dMRI) measure sensitive to diffusion within the intracellular (e.g. neuronal/axonal and glial) space, the Spherical Mean 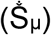; a quantitative MRI measure, the longitudinal relaxation rate R1 as a proxy for myelination; and a more widely-used general dMRI measure of ‘tract integrity’, the Fractional Anisotropy (FA), believed to reflect density of axonal packing, orientation of axons within a voxel and membrane permeability. Previous evidence suggests that changes to tuning functions within face-selective areas are linked to the development of face perception. Here, we used our data to test the hypothesis that, complementing the perceptual consequences of functional selectivity, microstructural changes in white matter tracts between areas in the face and body processing streams influence the development of facial expression and body posture perception, as well as the influence of body posture on facial expression perception.

## Methods

### Participants

A total of 45 typically-developing children (22 female) between 8 to 18 years of age (mean age = 12.96 ± 3.1) were recruited from the local community. The study took place over two visits and was part of a larger study focussing on brain development. Children underwent cognitive testing and a 3T MRI scan on their first visit, and a 7T scan (n = 44) on their second visit. All children had normal or corrected-to-normal vision and were screened to exclude major neurological disorders. All children had IQ values above 86 (Mean score ± SD = 107 ± 14.26, range = 86-145) (Wechsler Abbreviated Scale of Intelligence; Wechsler, 1999). Pubertal stage was determined using parental report on the Pubertal Development Scale (PDS) (Petersen et al., 1988). A strong positive correlation between age and PDS score was found (r_s_ = 0.85, p<0.0001). Therefore, in subsequent analyses, we used age adjustment only. Primary caregivers of children provided written informed consent prior to participating (with over-16s also providing their own written consent). Experimental protocols were approved by Cardiff University School of Psychology Ethics Committee and were in line with the Declaration of Helsinki. Participants were reimbursed with a voucher.

### Psychophysical testing

#### Stimuli

For our face stimuli, we used images of facial expressions from the Radboud and Karolinska Directed Emotional Faces validated sets of facial expressions (Langner et al., 2010; Lundqvist et al., 1998). Angry and disgusted facial expressions of four male identities were morphed separately using FantaMorph [FantaMorph Pro, Version 5]. The morphs changed in increments of 5%, leading to a total of 21 morph levels from fully angry to fully disgusted face generated for each identity (Figure 1).

**Figure 1.**
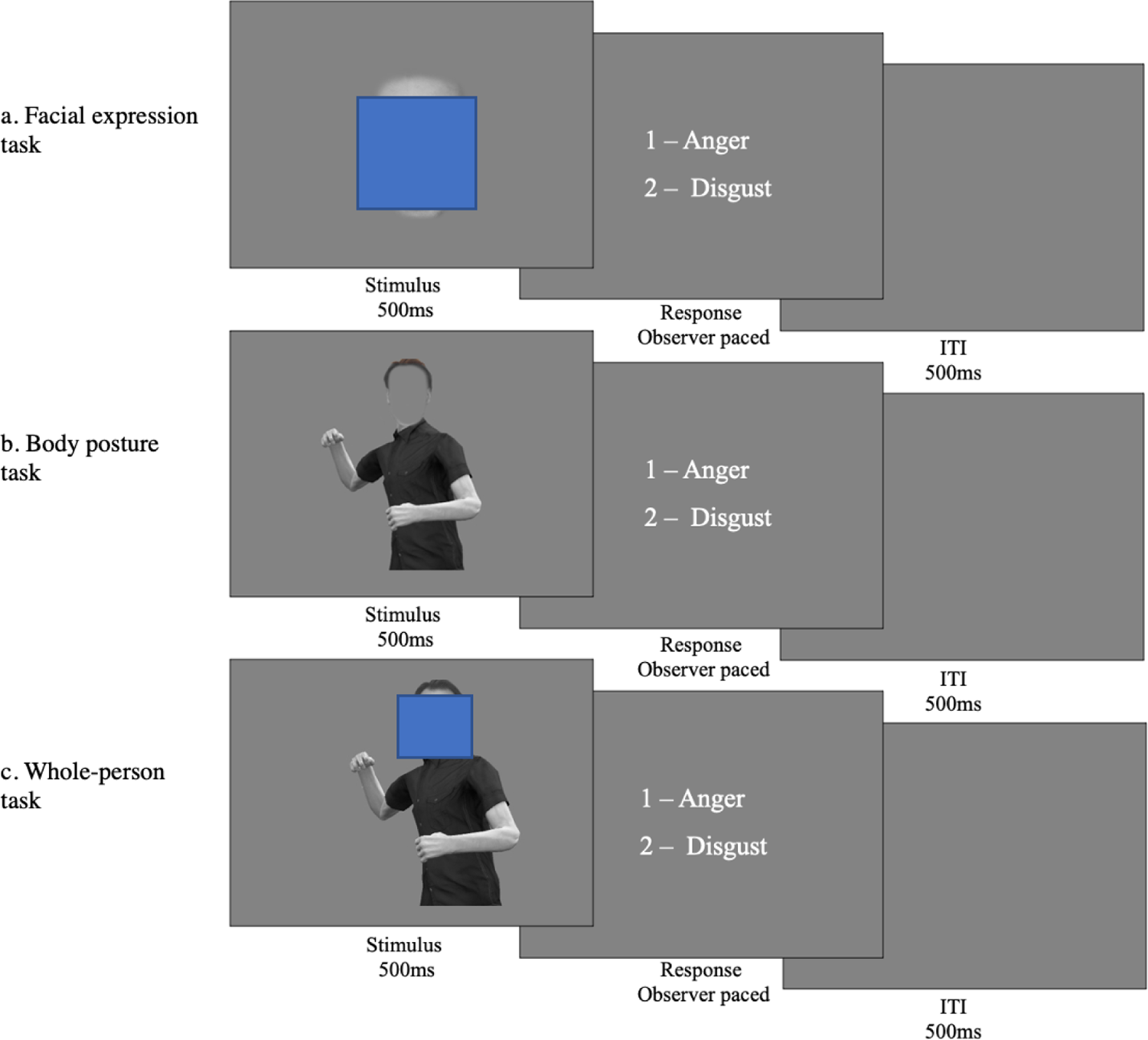
Psychophysical task design. The figure illustrates one trial for each of the three experimental tasks. The Facial Expression (A) and the Body Posture tasks (B) were used to index the precision of facial expression and body posture representations, respectively. The Whole-person task was used to measure the influence of body posture on facial expression perception. In the Facial Expression (A) and the Body Posture tasks (B) isolated faces or bodies were presented and participants were asked to categorize them as angry or disgusted. In the Whole-Person task (C), they were asked to base their categorization on the face and ignore the body. On each trial the stimulus was presented centrally for 500ms. Following presentation of the stimuli, response options appeared on screen and remained until an observer made a response. The next trial commenced following a 500ms inter-trial interval. Images of faces have been covered for this Preprint as BioRxiv does not allow them.

By contrast to facial expressions, body postures have to be morphed in 3D, rather than 2D image space. To create these stimuli, an actor dynamically posed emotional body postures while 3D location information was recorded from 22 body joints using a motion capture suit with motion trackers (Fedorov et al., 2018). The actors’ postures were modelled on typical body postures from the literature [e.g., Aviezer et al, 2008]. Based on the dynamically recorded behaviour of the actor, we chose four slightly different angry and disgusted body postures to create four identities. Visualisation of these postures was achieved using Unity 3D game engine [Unity, 2018]. Each of these postures was given a different identity by varying body composition and clothing. For each body identity, morphs between anger and disgust were generated via weighted averages of corresponding joint angles of the recorded postures. Similar to face morphs, body morphs changed in increments of 5%, resulting in a total of 21 body morph levels per identity ranging from fully angry to fully disgusted. Each body posture was combined with a facial identity to create a photorealistic ‘whole-person’ stimulus. For the body posture categorisation task, a mean grey oval was placed centrally over the face to conceal the distinguishing features of the facial expression (Figure 1). For the whole-person task, facial expression morphs were presented on a fully angry or a fully disgusted body (Figure 1).

#### Stimulus Presentation

We indexed the precision of (i) facial expression and (ii) body posture representations as well as (iii) the influence of body posture on expression perception by estimating psychometric functions in three separate psychophysical tasks (see Fig. 1 for details). Each task consisted of a 1-Alternative-Forced-Choice (1AFC) paradigm, in which participants viewed a single stimulus and were asked to categorize it as anger or disgust. In the Facial Expression and the Body Posture tasks, the stimulus was an isolated face or body morph. In the Whole-Person task, face morphs were presented either in the context of fully angry or fully disgusted bodies, and participants were instructed to categorize the facial expression and ignore the body posture. To optimize testing efficiency, the morph level of the stimulus presented on each trial was determined by the Psi method (Kontsevich & Tyler, 1999), a Bayesian adaptive procedure. Stimuli were presented centrally on the screen for 500ms. The images subtended approximately 15° visual angle (vertically) by 10° visual angle (horizontally). Following stimulus presentation, response options appeared and remained on screen until a response was recorded via a button press. A small number of 8 to 10-year-old participants indicated their response verbally and the experimenter pressed the button. Following each response, there was a 500ms inter-trial interval prior to the next stimulus appearing.

Presentation of the tasks was controlled by custom-written MATLAB (Version 2016b, The MathWorks, Natwick, MI, USA) code using the Psychophysics Toolbox (Version 3.0.14) (Brainard, 1997; Kleiner et al., 2007; Pelli, 1997) and the Palamedes Toolbox (Kingdom & Prins, 2010). For the facial expression and body posture tasks, there were 100 trials. For the whole-person task there were 496 trials: half were facial expression morphs on a fully angry body, and the other half on a fully disgusted body posture. Each of the four identities were displayed an equal number of times in each task. The order of tasks was fully counterbalanced across participants and each task had a short training phase at the beginning. Together, all three tasks took approximately 30 minutes to complete.

#### Analysis

Using custom-written Matlab code with the Palamedes toolbox (Kingdom & Prins, 2010), psychometric functions (PF) based on a cumulative Gaussian were fitted to the data of each task for each observer to extract the slope and point-of-subjective equality parameters (PSE). Lapse rate was fixed at 0.03, guess rate was determined by the experimental procedure and was fixed at 0. Two PFs were fitted to the two conditions within the Whole-Person task: one PF for data when the facial expression morphs were presented on an angry body, and the other for the facial expression morphs on a disgusted body. Goodness-of-fit of the PFs was assessed visually by two independent researchers. A total of 12 children were excluded from the facial expression condition, 6 children from the body posture condition, and 17 children from the whole-person condition due to poor fitting of the PFs.

For the Facial Expression and Body Posture tasks, the slope parameter was the key measure of interest. It provides an index for the precision of the perceptual representations underlying performance, with steeper slopes indicating greater ability to distinguish between subtle differences in the morphs. For the Whole-Person task, the key measure was the difference in PSE between the two conditions, i.e., when the same face morphs were presented on a fully angry or fully disgusted body. The PSE is the point at which the observer is equally likely to perceive the stimulus as either disgust or anger. The PSE change between the two conditions in the Whole-Person task reflects the contextual influence of body posture on facial expression judgements.

### Neuroimaging

On the participant’s first visit, diffusion MRI (dMRI) data were acquired on a 3T Connectom scanner (Siemens Healthcare, Erlangen, Germany) with 300mT/m gradients and a 32-channel radiofrequency coil (Nova Medical, Wilmington, MA, USA). The dMRI data were acquired using a multi-shell diffusion-weighted EPI sequence with an anterior-to-posterior phase encoding direction (TR/TE=2600/48ms; resolution=2×2×2mm; 66 slices; b-values=0 (14 vols), 500; 1200 (30 directions), and 2400; 4000; 6000 (60 directions)s/mm^2^; TA=16min 14sec). One additional volume was acquired with a posterior-to-anterior phase encoding direction for the purpose of EPI distortion correction. T1-weighted anatomical images were acquired using a 3D Magnetization Prepared Rapid Gradient Echo (MP-RAGE) sequence (TR/TE=2300/2ms; resolution=1×1×1mm; 192 slices; TA=5min 32sec). On the second visit, participants had structural and functional scans on a 7T Siemens Magnetom (Siemens Healthcare, Erlangen, Germany). Whole-brain echo-planar imaging (EPI) gradient echo data were acquired (TR/TE=2000/30ms; resolution=1.5×1.5×1.5mm; 87 slices; multi-band factor=3; TA=9min 18sec), with slices angled along the anterior commissure and posterior commissure to minimise signal drop out from the temporal regions. In addition, a B0 field map (TR/TE 1/TE 2=560/5.1/6.12ms; resolution=3×3×3mm; 44 slices; TA=1min 7sec) was acquired to unwarp the high-field EPI data. A high-resolution MP2RAGE structural scan (TR/TE=6000/2.7ms; resolution =0.65×0.65×0.65mm; TA=10min 46secs) was also acquired (Marques et al., 2010). An SA2RAGE B1 map (TR/TE=2400/0.72ms; resolution=3.25×3.25×6mm; TA=1min 26sec) was acquired to bias-field correct the MP2RAGE (Eggenschwiler et al., 2012). Retrospective correction of head movement was used to mitigate blurring of the high-resolution MP2RAGE data at 7T (Gallichan et al., 2016).

For the functional localiser task, which was used to identify cortical regions involved in face and body processing for each individual, participants were presented with grey-scale images of faces, houses, bodies, and chairs. Images were shown in blocks by stimulus category, and the order was pseudorandomised across participants. Within each block 15 images were displayed for 800ms each with a 200ms inter-stimulus interval. A block of fixation followed the stimulus category block, and the participant was asked to focus on the fixation cross displayed centrally on a mean grey screen (15s). For the last second of each fixation block, the fixation cross turned red to indicate that the next block of images would commence. In total there were four blocks of each stimulus category. Therefore, for each stimulus category, 64 trials per condition were presented overall. To ensure participants attended to the images, they were instructed to respond using a key press on an MR-compatible button box if the same image was presented twice in succession (1-back task). The number of repeated images per block varied between 0 and 3. The average accuracy on the 1-back task was 90% (SD ± 0.18), indicating a high degree of attention to the stimuli. Participants completed a short practice version of the task prior to the scan.

#### Functional MRI processing

Functional MRI data processing was carried out using FEAT (FMRI Expert Analysis Tool, Version 6.0, FMRIB Functional Magnetic Resonance Imaging of the Brain Software Library FSL; (Jenkinson et al., 2012). High-field functional data was unwarped using the B0 field map generated from a brain extracted magnitude and phase image in FSL (Woolrich et al., 2001). The 7T functional data was then registered into 3T diffusion space to allow for tractography in native diffusion space. First, the pre-processed high-resolution MP2RAGE (skull stripped, bias corrected, neck cropped) image was registered to the diffusion data using Advanced Normalization Tools (ANTs) (Avants et al., 2011). The T1w contrast from MP2RAGE was selected as the best intermediary between functional and diffusion data sets. Next, 7T functional data were registered to the diffusion-aligned T1w image. In addition, functional data were transformed to Montreal Neurological Institute (MNI) space using FMRIB’s Linear Image Registration Tool (FLIRT) (Jenkinson et al., 2002; Jenkinson & Smith, 2001).

Motion correction of fMRI data was performed with FSL’s Motion Correction using FMRIB’s Linear Image Registration Tool (MCFLIRT); one participant was removed from the analysis due to motion of more than 2 voxels (3mm) during the fMRI task. The average absolute motion for all participants was 0.74mm ± 0.7. To mitigate the effects of motion in the functional imaging analysis, the estimated motion traces from MCFLIRT were added to the GLM as nuisance regressors (Jenkinson et al., 2002). Skull-stripping and removal of non-brain tissue was completed using Brain Extraction Tool (BET; (Smith, 2002). All data were high-pass temporal filtered with a gaussian-weighted least-squares straight line fitting (sigma=50.0s). On an individual subject basis, spatial smoothing was applied using a Gaussian smoothing kernel with a Full Width at Half Maximum (FWHM) 4mm to preserve high spatial resolution (Woolrich et al., 2001). Time-series statistical analysis was carried out using FMRIB’s Improved Linear Model (FILM) with local autocorrelation correction (Woolrich et al., 2001).

A univariate GLM was implemented to examine the BOLD response associated with face and body stimuli. The contrast Faces > Houses as well as the contrast Bodies > Chairs were used to localise cortical regions involved in face and body processing, respectively. Functionally-defined regions-of-interest (ROIs) involved in face processing (FFA, OFA and pSTS-face) and body processing (FBA, EBA, pSTS-body) were identified on a subject-by-subject basis in individual subject space. Z-statistic images were uncorrected and thresholded at p=0.1. ROIs could be identified in the following number of subjects: OFA (n=37); FFA (n=43); pSTS-face (n=41); EBA (n=39); FBA (n=40); pSTS-body (n=39). ROI analyses (and therefore all subsequent white matter tractography and analyses) were restricted to the right hemisphere, as too few ROIs could reliably be identified in the left hemisphere across participants. As a validation step, the individual ROIs were translated into MNI space, and an average coordinate across participants was determined for each ROI [Table 1]. The Euclidean distance was calculated between each ROI and the average coordinates in MNI space for each participant to provide a measure of how variable the locations were across participants. The average coordinates were comparable to other studies where these regions have been extracted in MNI space (Bona et al., 2015; Harry et al., 2016; Schobert et al., 2018; Vocks et al., 2010). ROIs were inflated to 10mm in diameter into surrounding WM to be used as seed regions for functionally-defined white matter tractography. In addition, an ROI in the right ATL was manually drawn based on anatomical landmarks as follows: a coronal plane in right temporal lobe extending from the lateral fissure to the ventral surface of the brain (Hodgetts et al., 2015), with the position of the plane just anterior to where the central sulcus meets the lateral fissure.

**Table 1.** Average coordinates for ROIs. The coordinates reported are the average coordinates of all individual participant ROIs registered to MNI space, in order to allow for comparison with coordinates reported in the literature. The Euclidean distance was calculated between each ROI and the group average coordinate in MNI space, for each observer. The standard deviation (SD) of the Euclidian distance between individual ROIs and group average coordinates is reported to provide a measure of how variable the ROIs were across observers.

| **ROI** | **x** | **y** | **z** | **SD** |
| --- | --- | --- | --- | --- |
| **FFA** | 38 | -43 | -20 | 7.1 |
| **OFA** | 38 | -76 | -7 | 11.24 |
| **pSTS-face** | 46 | -57 | 12 | 6.04 |
| **FBA** | 38 | -43 | -18 | 7.6 |
| **EBA** | 41 | -78 | 0 | 9.17 |
| **pSTS-body** | 44 | -59 | 11 | 8.87 |

#### Diffusion-Weighted-Imaging pre-processing

Diffusion-Weighted-Imaging (DWI) data were pre-processed to reduce thermal noise and image artefacts which included image denoising (Veraart et al., 2016), correction for signal drift (Vos et al., 2017), motion, eddy current, and susceptibility-induced distortion correction (Andersson & Sotiropoulos, 2016), gradient non-linearities, and Gibbs ringing (Kellner et al., 2016). The pre-processing pipeline was implemented in MATLAB, but depended on open-source software packages from MRtrix (Tournier et al., 2019) and FSL (Jenkinson et al., 2012). DWI data quality assurance was performed on the raw diffusion volumes using slicewise outlier detection (SOLID) (Sairanen et al., 2018). Digital brain extraction was performed using the FSL Brain Extraction Tool (BET) (Jenkinson et al., 2012) followed by the FSL segmentation tool (FAST) which segmented tissue into CSF, WM and GM.

#### Tractography

In order to identify tracts, we first obtained voxel-wise estimates of fibre orientation distribution functions (fODFs). This was done by applying multi-shell multi-tissue constrained spherical harmonic deconvolution (Jeurissen et al., 2014) to the pre-processed images (Descoteaux et al., 2009; Seunarine & Alexander, 2014; Tournier et al., 2004, 2007) with maximal spherical harmonics order lmax = 8. Functionally-defined fibre tracts (FDFTs) were generated using the using the b=6000s/mm^2^ shell for each participant between ROIs in the right hemisphere within the face network (OFA to FFA (Figure 2), FFA to ATL, pSTS-face to ATL) and the body network (EBA to FBA) in each subject. In order to assess the confluence of face-and body-selective networks in ATL, we also identified FDFTs between FBA to ATL and pSTS-body to ATL. Streamlines were generated using a probabilistic algorithm in MRtrix using one ROI as seeding mask and the second as an inclusion region, following the organisation of the visual processing hierarchy. No streamlines were found between OFA/FFA and pSTS-face, nor between EBA/FBA and pSTS-body. All identified FDFTs were visually inspected, and spurious fibres manually removed. Any FDFTs between ROIs with less than 20 streamlines were removed from subsequent analyses (FFA-ATL: n=1, pSTS-face-ATL: n=1, FBA-ATL: n=1, EBA-FBA: n=1). In addition to the functionally-defined fibre tracts generated, two anatomically-defined tracts, as comparison tracts, were extracted in the right hemisphere: the inferior longitudinal fasciculus (ILF) and the cortico-spinal tract (CST). ILF was selected as a comparison tract to reconstructed FDFTs as it traverses similar subcortical regions, therefore providing an index of specificity, and CST was selected as a region outside the visual processing regions. TractSeg segmentation software (Wasserthal et al., 2019) was used to automatically extract these tracts.

**Figure 2.**
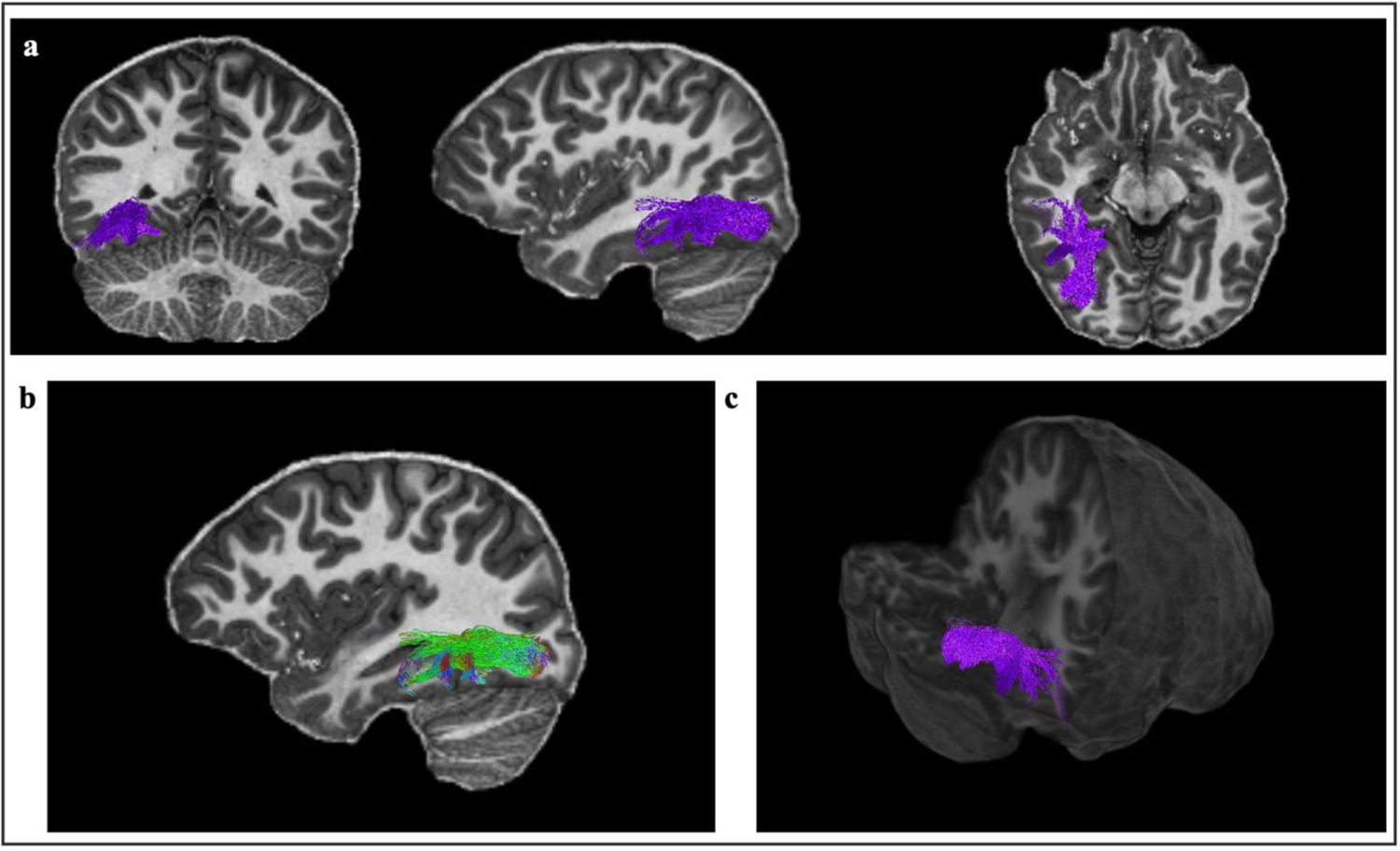
Reconstructed FDFTs for OFA-FFA for one participant. The top panel depicts the tracts in a coronal, sagittal and axial planes [a]. The tracts are shown with directional colour encoding in the sagittal plane [b], where green is anterior-posterior, red is left-right and blue is superior-inferior. A 3D positioning of the tracts is shown in the bottom right panel [c].

The following parametric maps were computed to characterise the microstructure of each fibre tract. Diffusion-weighted data acquired across 3 b-value shells (b=500, 1200, 2400s/mm^2^) were fitted to a cumulant expansion of the signal up to the 4th order cumulant (i.e., kurtosis), to improve estimation of the 2nd order cumulant (the diffusion tensor), and its derived measures (e.g., FA) (Jensen et al., 2016; Veraart et al., 2011). FA has been widely interpreted in the literature as a measure of ‘tract integrity’, but is actually thought to reflect multiple aspects of the microstructure, including density of axonal packing, orientation of axons within a voxel and membrane permeability (Jones et al., 2013). While FA is more general in its interpretation, it is a powerful benchmark for comparison with existing cohort studies of development (Lebel et al., 2019; Lebel & Beaulieu, 2011; Tamnes et al., 2018). We complemented FA with a second diffusion measure, the spherical mean 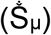 of the diffusion-weighted signal at b=6000 s/mm2, which has been shown to suppress extracellular signal (Kaden et al., 2016; Veraart et al., 2011). The resulting signal reflects restricted diffusion, i.e., water that is trapped intracellularly, such as in axons of neurons, or glial cells, without having to rely on signal modelling. In this work, 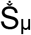 was used to refine the inferences from FA, as an increase in 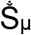 is more directly influenced by intracellular signal, such as increases in the number or density of neuronal and glial processes (versus the surrounding microstructure at large) than FA. The third measure used was derived from quantitative MRI, the longitudinal relaxation rate R1 (= 1/T1), extracted from the MP2RAGE sequence (Marques et al., 2010). R1 is sensitive to myelination and has been shown to correspond with white matter maturation across childhood and adolescence (Lutti et al., 2014). Diffusion and relaxometry measures were projected across streamlines and averaged for each tract resulting in one average metric value per FDFT or comparison tract in each participant.

### Statistical analysis

All statistical analyses were run in R (Version 3.6.1). To assess the relationship between behavioural metrics and FDFTs, Spearman correlations were performed as data were non-normally distributed. Bonferroni correction was used to control for multiple comparisons of the six FDFTs with a corrected value of p<0.008. Age-related variability in significant tract-behaviour relationships was controlled for using multiple linear regression. As this was a secondary analysis, the significance level was set at p<0.05.

## Results

### Behavioural results

In line with previous research (Dalrymple et al., 2017; Herba et al., 2006), our data indicate that facial expression discrimination improves with age in children and throughout adolescence, as indicated by a significant positive relationship between facial expression precision and age (r_s_= 0.51, p<0.005) [Figure 3]. While not reaching statistical significance, a positive relationship was observed between body posture discrimination ability and age (r_s_= 0.27, p=0.078) [Figure 3], suggesting that body posture discrimination might also improve with age.

**Figure 3.**
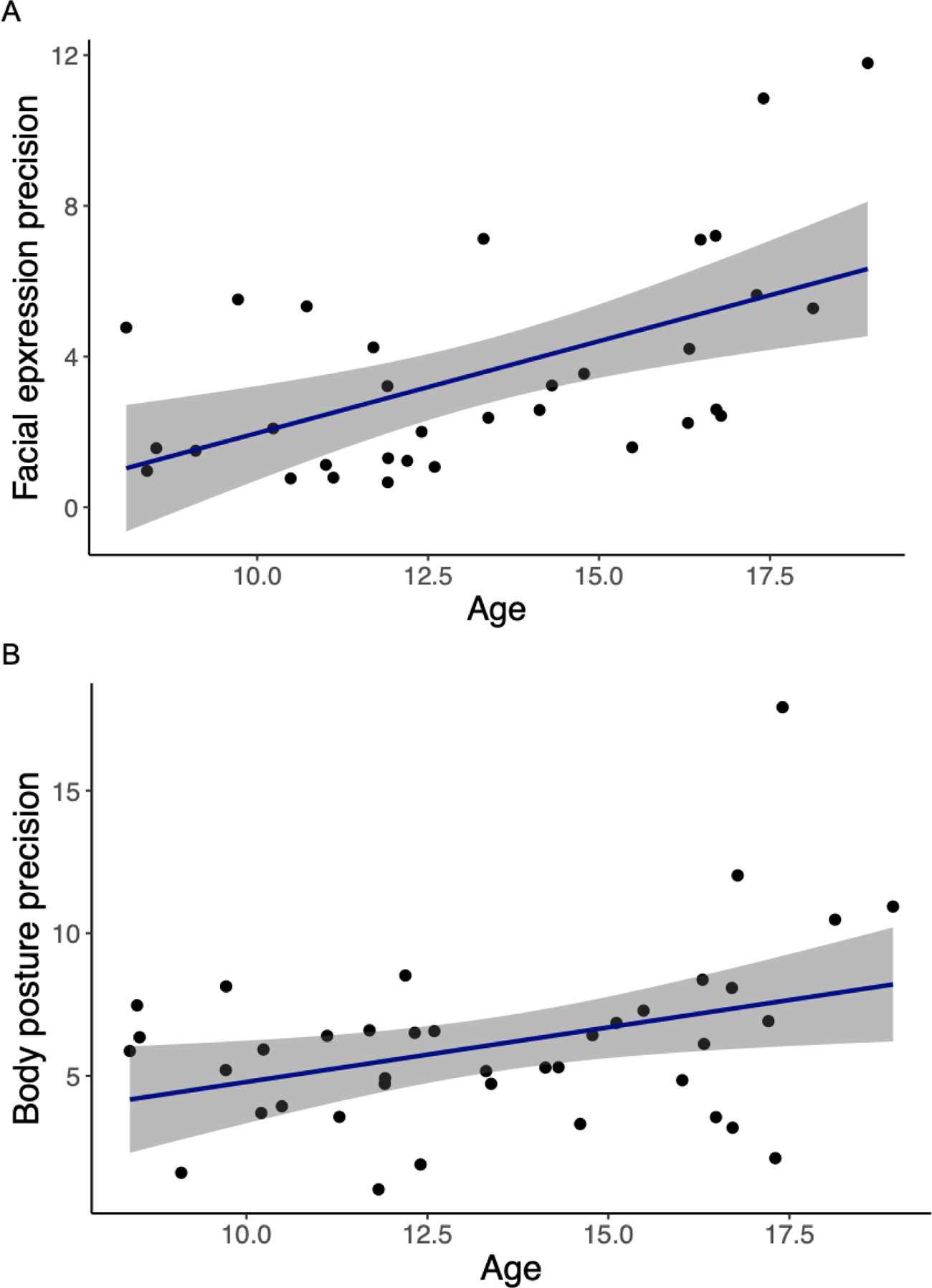
Relationship between facial expression precision and body posture precision with age. The correlation between age (years) and (A) facial expression precision (r_s_= 0.51, p<0.005), and (B) body posture precision (r_s_= 0.27, p=0.078), as indexed by the slope estimate of the individual’s PF for each task. Each point represents one observer. The 95% confidence interval is shown with grey shading.

As expected, body posture clearly influenced facial expression perception in our participants [Figure 4]. Specifically, perception of facial expressions was biased towards the body emotion, as indicated by a robust difference in the PSE when the same facial expressions were presented in the context of a fully angry vs. a fully disgusted body posture (t(28)= 5.10, p<0.0001). Interestingly, the influence of body posture on facial expression perception decreased with age in children and throughout adolescence, as indicated by a negative relationship between PSE change and age (r_s_ = −0.39, p=0.0395) [Figure 5]. In other words, younger children were more strongly influenced by body posture in their categorisation of facial expressions than older children and adolescents. There was no significant relationship between facial expression precision and PSE change (r_s_ = −0.32, p=0.11; controlling for age r_s_ = −0.32, p=0.12), neither was there a significant relationship between body posture precision and PSE change (r_s_= −0.13, p=0.55; controlling for age r_s_ = −0.13, p=0.57). Controlling for facial expression precision in the relationship between PSE change and age resulted in the relationship no longer being significant (r_s_= −0.29, p=0.16), whereas controlling for body posture precision in the relationship between PSE change and age had no effect on the relationship (r_s_ =-0.44, p=0.03). Together, these findings suggest that the improvement in facial expression precision is a key factor driving the reduction in body context influence with age, whereas improved body posture discrimination does not have an effect on this relationship.

**Figure 4.**
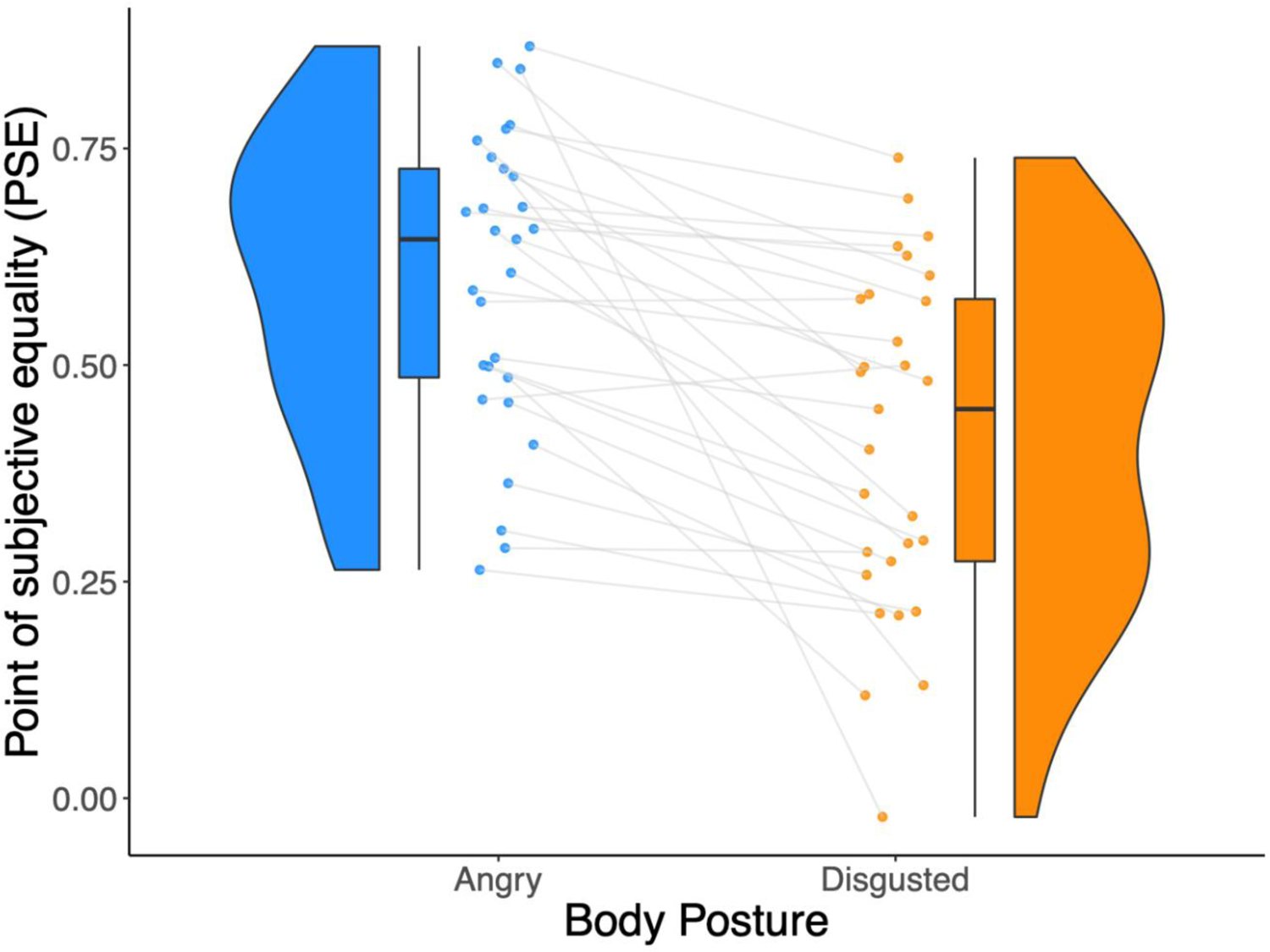
PSE change between categorisation of facial expressions on a 100% angry and a 100% disgusted body posture. The raincloud plot displays the PSE values for the whole-person condition when facial expressions were presented with a fully-angry (blue) or fully-disgusted (orange) body posture. A significant difference (t(28)= 5.10, p<0.0001) was observed between the PSE values when the facial expression was categorised with a 100% angry and 100% disgusted body posture. Each line represents one observer and depicts the change in PSE. The distribution of the values is illustrated by the shaded area, with the boxplot indicating the median and the interquartile range.

**Figure 5.**
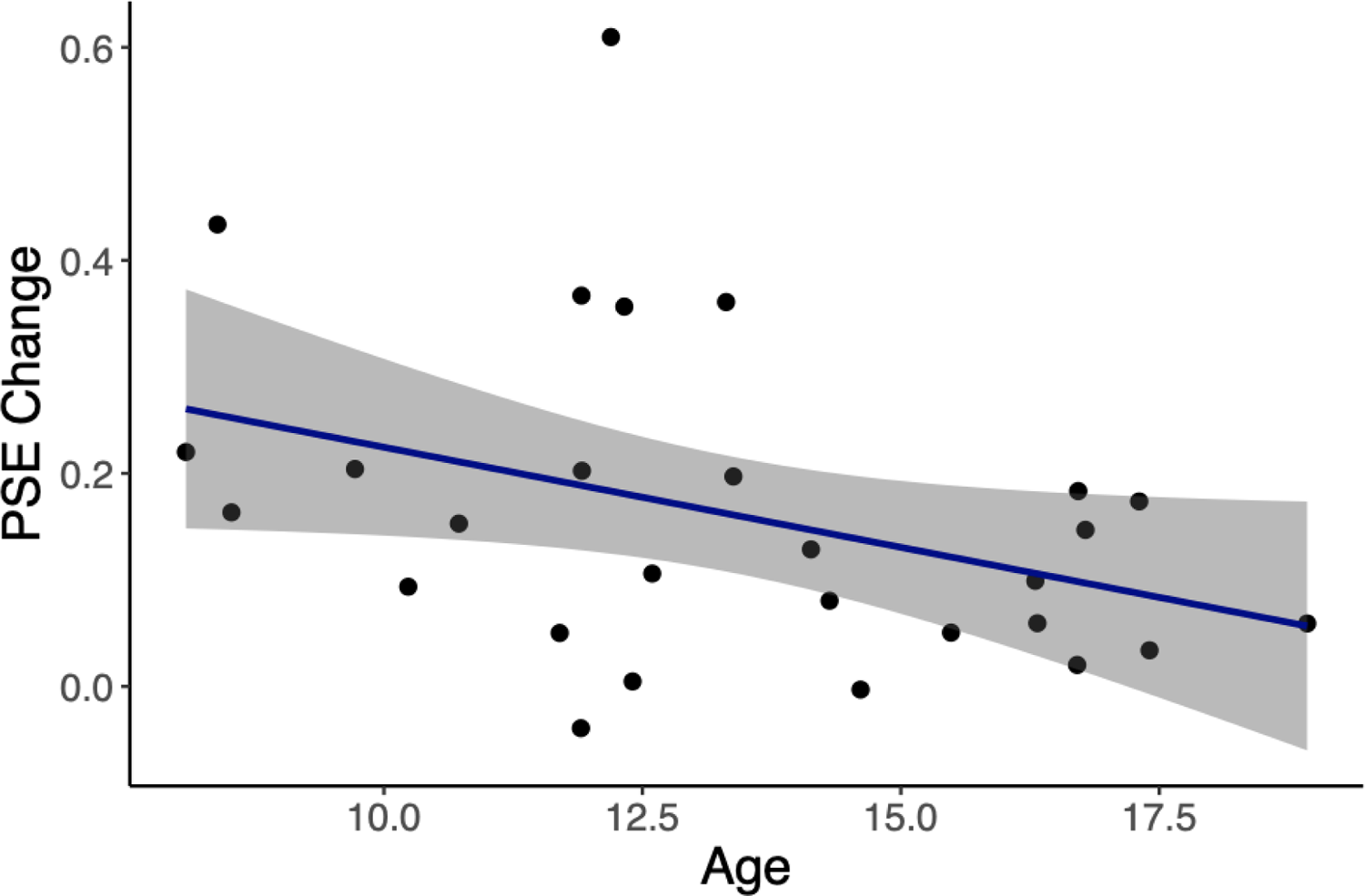
Relationship between PSE change and age. Significant negative correlation between age (years) and PSE change (r_s_ = −0.39, p=0.0395). Each point represents one observer. The 95% confidence interval is shown with grey shading.

### Tractography results

In order to isolate different aspects of white matter microstructure across development and their relation to perception, we used 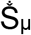 to index characteristics directly linked to diffusion within the intracellular (axonal, glial) space, R1 as a measure of myelination, and FA as a widely-used general measure reflecting multiple aspects of microstructure. As described below, these measures related differentially to age, perception, as well as the interaction between age and perception.

#### Age and microstructural change

Changes to intraaxonal and/or glial signal fraction, 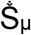, within all FDFTs of the face and body networks (with the exception of FBA to ATL tracts) were observed across development, as indicated by a positive relationship between 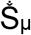 and age (all p’s<0.005). Similarly, the index of myelination (R1) of all face-and body-related FDFTs increased from 8 to 18 years, as demonstrated by the relationship between R1 and age (all p’s<0.005). There were no significant relationships between the more general FA measure in any FDFTs and age. In addition to FDFTs, R1_ILF_ was significantly positively correlated with age (r_s_= 0.65, p < 0.001), as was 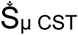 (r_s_= 0.61, p < 0.0001) and 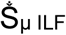 (r_s_= 0.56, p= 0.0001).

#### Perceptual performance and microstructure

Better facial expression discrimination was related to an increase in all three microstructural measures for the tracts linking OFA and FFA (FA_OFA-FFA_: r_s_= 0.55, p=0.006; Figure 6; 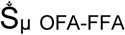: r_s_= 0.55, p=0.007; R1_OFA-FFA_: r_s_= 0.62, p=0.001). No other significant relationships were observed between facial expression precision and FDFT measures within the face and body networks. More accurate body posture perception was associated with changes to intracellular signal, 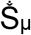, and increased myelination (R1) of the tracts linking OFA and FFA, as indicated by a positive relationship between perception and both 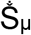 (r_s_= 0.60, p=0.0009) and R1 (r_s_= 0.59, p=0.001). No other significant relationships were found between body posture precision and FDFT measures, including FA, across all remaining tracts. Finally, a greater influence of body context on facial expression perception was related to reduced 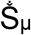 (r_s_= −0.61 p=0.002), as well as reduced R1 (r_s_= −0.61 p=0.002) in tracts linking pSTS_body_ and ATL.

**Figure 6.**
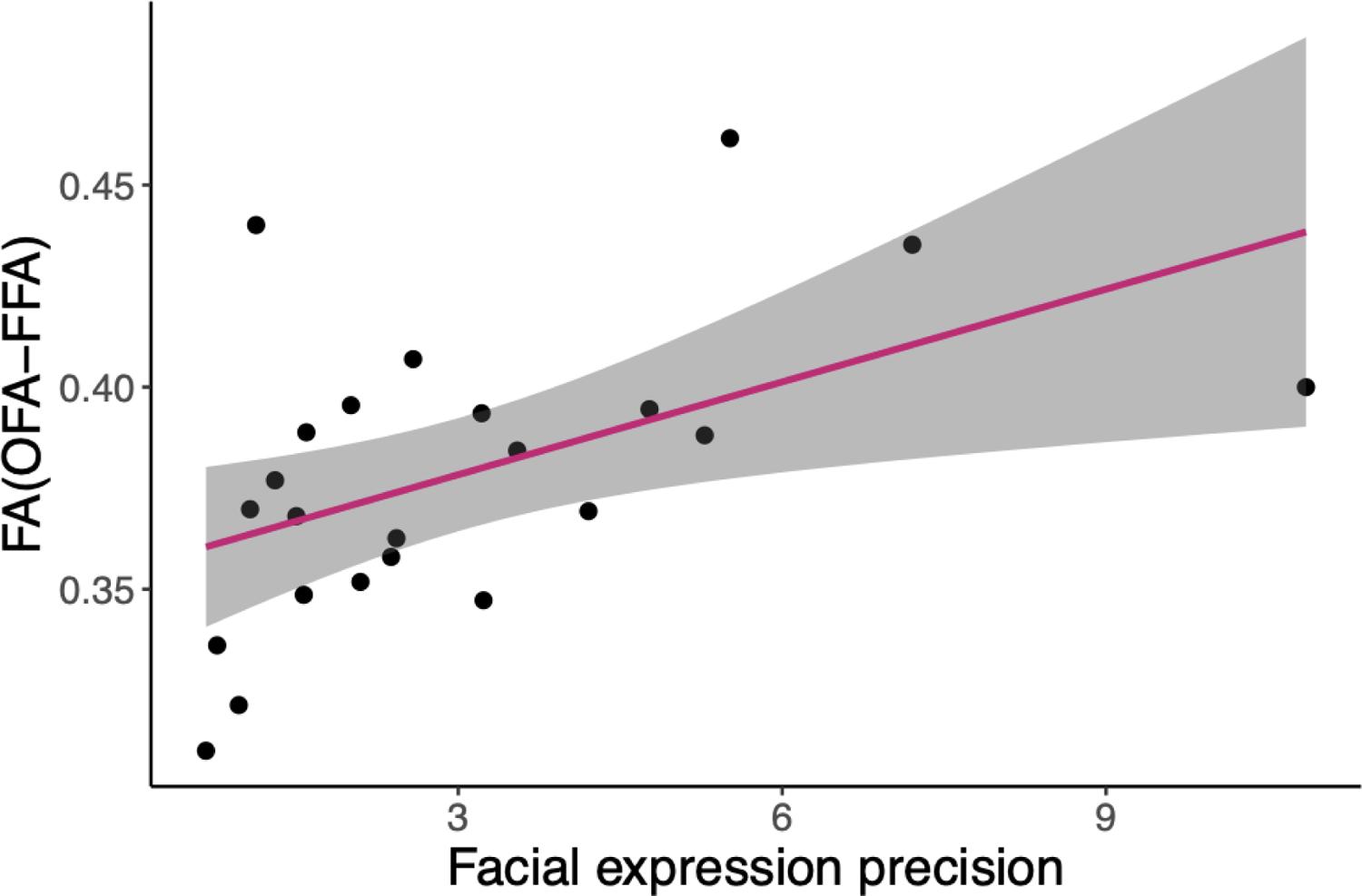
Relationship between facial expression precision and FA_(OFA-FFA)_. There was a significant correlation between facial expression precision and FA of OFA-FFA tracts (r_s_= 0.5526, p=0.006). Each point represents one observer. The 95% confidence interval is shown with grey shading.

To explore the specificity of these findings to FDFTs within face and body networks, ILF tract metrics were assessed in relation to the behavioural measures, and a significant relationship was found between R1_ILF_ and PSE change (r_s_= −0.59, p=0.002). No other significant relationships were found. A comparison tract outside of visual processing regions, the CST, also showed no significant relationships between microstructural measures and behaviour.

#### Age, perceptual performance, and microstructural change

To identify more precisely which microstructural aspects were related to perceptual performance compared to more general age-related brain development, we conducted an analysis controlling for the effect of age on tract microstructure when predicting perception. Specifically, we first implemented a linear regression model using age to predict tract microstructure and then used the residuals of this analysis in a second regression model as a predictor for perception.

Once age-related changes were controlled for, facial expression perception was differentially associated with microstructural characteristics of the FDFT linking OFA and FFA. Specifically, the residuals of the age-microstructure analysis for FA_OFA-FFA_ predicted facial expression precision [F(1,21) = 4.34, p=0.050], suggesting that the microstructural characteristics of the OFA-FFA tracts captured by FA were associated with the ability to recognise facial expressions independently of age-related brain development. Equivalent regression analyses using the age-microstructure residuals of 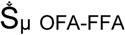 [F(1,21) = 1.09, p=0.31] or R1_OFA-FFA_ [F(1,21) = 0.15, p=0.71] were not significant, suggesting that these tract measures no longer predicted facial expression precision after accounting for age-related changes. Together, these analyses suggest that general age-related changes drive the relationship between facial expression perception and 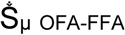 and R1_OFA-FFA_. By contrast, our data suggest that FA _OFA-FFA_ captures microstructural characteristics that are related to perception independently of age.

The relationship between body posture perception and microstructural measures of the OFA-FFA tracts appeared to be mainly driven by general age-related brain development only. Specifically, the residuals of the age-microstructure analysis for Š_µ OFA-FFA_ [F(1,26) = 2.50, p=0.13] and R1 _OFA-FFA_ [F(1,26) =2.25, p=0.15] were not related to body posture perception once age-related microstructural changes were controlled for, suggesting that age accounted for most of the relationship between these metrics and body posture perception. The relationship between R1_ILF_ and PSE change also appeared to be mainly driven by age, as the residuals of the age-microstructure regression [F(1,23) = 0.27, p = 0.61] were not related to PSE change after accounting for age-related variance.

Finally, our data suggest a similar situation with respect to the relationship between microstructural characteristics of the tract linking pSTS_body_ with ATL and the influence of body context on facial expression perception. The residuals of the age-microstructure analysis for 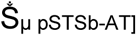 [F(1,22) = 0.22, p=0.64] and R1_pSTSb-ATL_ [F(1,22) = 0.23, p=0.63] no longer related to PSE change once age-related changes in these metrics were controlled for.

## Discussion

In the current study, we measured facial expression and body posture perception in children and adolescents from 8 to 18 years, as well as the influence of body context on facial expression perception. We directly linked perceptual performance to microstructural changes within white matter tracts along face-and body-selective processing pathways. We show that, using multiple, complementary microstructural measures, we can identify both age-related developmental changes in microstructure related to behaviour, as well as age-independent microstructural changes. 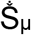 and R1, specific measures based on intracellular diffusion-weighted signal and myelination respectively, increased with age along face-and body-selective pathways. This increase was not seen in FA, a more general, composite measure that indexes several aspects of the microstructure. Increases in all three measures in the tracts linking OFA and FFA related to developmental improvements in the ability to recognise facial expressions. Importantly, however, once microstructural changes due to age were controlled for, only the composite measure FA of the OFA-FFA tracts predicted perceptual performance, while the other two measures were not related to behaviour independently of age. In addition to improvements in facial expression perception with age, we found that children and adolescents are less influenced by body posture when perceiving facial expressions as they get older. Our data suggest that this decreasing influence of body posture is driven by improvements in facial expression discrimination ability. The influence of body context on facial expression perception was related to microstructural measures based on intracellular diffusion-weighted signal and myelination, 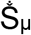 and R1 respectively, of the tract between pSTS_body_ and ATL. However, this relationship was mainly linked to age-related development of this tract rather than being driven by perception independently of age. Taken together, our results show that behavioural variability across development is driven by both age-related maturation of white matter, as well as age-independent variation in structural properties which contribute to behaviour.

The importance of functional characteristics of face-selective regions for social perception has been well-studied in adults, and, to a lesser extent, in children and adolescents. Specifically, development of the ability to recognize facial identity is linked to increases in the size of face-selective FFA, as well as increased response amplitude and selectivity to faces in this region (Golarai et al., 2009, 2010; Gomez et al., 2017; Natu et al., 2016). However, the role of white matter connections along the processing hierarchy in constraining function and behaviour is less well-understood. In adults, the properties of tracts at the boundary between white and grey matter close to FFA have been linked to face recognition ability (Gomez et al., 2015; S. Song et al., 2015). Here, we show that the structure of tracts linking OFA to FFA contributes to facial expression recognition ability during development. This relationship was specific to tracts linking face-selective regions and was not found for tracts linking body-selective EBA and FBA, despite the latter being in close proximity to FFA. Similarly, the absence of a relationship between facial expression discrimination ability and microstructural characteristics of large occipitotemporal association tracts like the ILF suggests behavioural specificity exists within smaller fibre bundles linking face-specific regions. This finding provides support for previous suggestions that tracts linking face-specific regions along VTC are distinct from ILF (Gomez et al., 2015; Wang et al., 2020).

Microstructural MRI measurements are made at the level of imaging voxels (mm scale), but inferences are drawn at the level of the microstructure within that voxel (µm). This situation can lead to a lack of specificity in identifying those elements of the microstructure that are underlying changes in the observed metrics. Our aim was to address this issue by using multiple microstructural measures within the same sample, which are optimised to detect signals arising from different components of the microstructure. In particular, the ultra-strong gradients of the Connectom scanner (300mT/m) allowed us to obtain diffusion data at high b-values (b=6000s/mm^2^) while maintaining a reasonable signal to noise ratio (SNR). At these b-values, the signal is weighted more heavily towards the diffusion of water molecules in the intracellular space, providing the rotationally invariant spherical mean 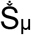 metric with greater sensitivity to the intracellular components of the microstructure. An increased 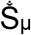 measure could, for instance, reflect reduced axon diameter or a greater number and/or density of neuronal and/or glial processes. By contrast, the fractional anisotropy (FA) measure, which is arguably the most widely-used measure of white matter microstructure from diffusion MR, has high sensitivity in detecting a range of changes in microstructure, however it is not possible to disentangle the microstructural components driving these changes (Afzali et al., 2021; Jones et al., 2013). FA reflects the (normalised) standard deviation of diffusivities along three orthogonal axes: the principal diffusion axis, and the two orthogonal axes. When diffusion is the same along all three axes, FA is zero. When constrained to move along just one axis, the FA assumes a value of one. FA can therefore reflect multiple aspects of the microstructure, including orientation of axons within a voxel, changes in density of axonal packing, membrane permeability, as well as combinations of these factors. In order to address myelination more directly, given its important role in development of white matter, we also obtained measures of the longitudinal relaxation rate (R1), which has previously been shown through comparative ex-vivo studies of MR measurements and histology to map myelin content closely (Lutti et al., 2014).

What can these measures tell us about the link between behaviour and white matter microstructure during development? Our findings suggest that the more specific measures of 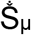 and R1, sensitive to the signal from intracellular space and myelin, respectively, are strongly age dependent. These measures increased with age across almost all of the tracts that we studied here. This finding is consistent with previous research, which found that 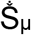 is sensitive to age-related maturation of white matter across the developing brain (Raven et al., 2020), with age-related changes in 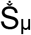 suggested to reflect reduced axonal diameter, greater complexity of neuronal and glial processes, and increased myelination associated with more advanced developmental stage. Similarly, R1 has previously been shown to increase with age across the large fascicles in the brain (Yeatman et al., 2014). The positive relationship between facial expression discrimination ability and 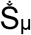 and R1 of tracts linking OFA to FFA suggests that changes in number, density and/or complexity of neuronal and glial processes with general development, as well as myelination levels, contribute to the improvement in facial expression discrimination performance with age. Conversely, the FA measure was relatively insensitive to age across all tracts identified here and was predictive of facial expression discrimination ability even after removing age-related variability. These results suggest that there may be aspects of microstructure that explain behavioural variability beyond age-related changes in neuronal and glial processes and myelination. While caution is warranted in interpreting this finding, given that we cannot specifically identify what these structural changes might be, previous research in adults can provide some insight. Specifically, variability in FA linked to facial identity processing ability in adults in white matter close to FFA has been proposed to be due to an increase in the number and density of connections to neurons in FFA (Gomez et al, 2015). Support for a similar proposal applying to the current data is also realised in recent findings reporting large variability in FA along tracts during development, which dwarf any age-related changes in FA across tracts (Yeatman et al, 2014). Whether our findings of age-dependent and age-independent microstructural variability reflect a general organising principle for the constraints posed by white matter structure on behaviour during development, or whether this is specific to the structure-behaviour relationship within this particular network remains an open question. In order to disentangle the characteristics of white matter structure driving behavioural changes during development and those underpinning naturally occurring behavioural differences within the population, it will be important for future research to directly compare multiple microstructural measures and their contribution to perceptual ability across children and adults.

Although white matter tracts linking regions of the face perception network have previously been identified (Gschwind et al, 2012; Pyles et al, 2013; Wang et al, 2020), much less is known about white matter connections linking body-specific cortical regions. To our knowledge, our study was the first to use body posture morphs to investigate the development of body posture discrimination ability. The results indicate that body posture perception follows a similarly protracted developmental trajectory to facial expression perception, reaching far into adolescence. To relate this perceptual development to brain structure, we mirrored the anatomical and hierarchical processing of the face network in our extraction of tracts for the body network. Interestingly, we found no evidence for a relationship between body posture discrimination ability and the structure of tracts linking EBA to FBA, but instead found evidence for a relationship with the structure of tracts linking OFA to FFA. This relationship was limited to the 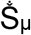 and R1 measures however, and was not significant after removing age-related variability. This finding suggests that variability in these tracts related to body posture discrimination appears to be limited to age-related maturation of white matter. The notion that tracts linking face-specific processing regions also encode elements of body posture discrimination is consistent with previous research showing similarly robust levels of activation of FFA in response to blurred faces on bodies as to faces alone (Cox, 2004). Nevertheless, it is still surprising that we do not find an association between body posture perception and characteristics of tracts linking body-selective areas, while we do find this link for face-selective areas. Importantly, the relationship between facial expression discrimination ability and the microstructure of OFA-FFA tracts remained significant even after removing variability associated with body posture discrimination, highlighting that facial expression perception ability is related to tract microstructure independently of body posture perception ability. The lack of significant associations found between body posture perception and tracts linking body-specific regions may reflect our gaps in knowledge regarding white matter connections and pathways important in body perception.

A key aim of our study was to identify not only the relationship between brain development and perception of isolated faces and bodies, but also its relation to perception of *integrated* facial expression and body posture signals. In everyday life, facial expressions are seen within a context of other socially-relevant signals, most importantly, the other person’s body posture. It is well-established that body context has an important influence on facial expression perception in adults, and that this effect is larger in children. Our data suggest that the influence of body context on facial expression perception continues to decrease well into adolescence. Moreover, our results point towards a potential mechanism for the decreased reliance on body posture with age. Specifically, the data suggest that the improvement in facial expression discrimination ability drives the decrease in the body context effect across development. It is tempting to speculate that a similar mechanism might drive differences observed throughout the lifespan. For instance, older adults rely more strongly on body context in facial expression perception relative to younger adults (Abo Foul et al., 2018; Kumfor et al., 2018). Given that older adults also suffer a reduced ability to discriminate between different facial expressions, it is possible that the increased reliance on body posture is a direct result of their reduced facial expression discrimination ability, a mirror-analogue to what we found here in children and adolescents.

It is important to highlight that our results cannot be explained by children simply ignoring the instructions and judging body context instead of facial expression in the whole-person task. The high consistency of performance within participants in discriminating facial expressions, as measured by the slope of the PF, across the conditions with and without body context (r_s_= 0.36, p=0.07) clearly indicates that children followed task instructions and discriminated facial expressions in the whole-person task, rather than simply judging the body posture. Additionally, based on our exclusion criteria, any participants, who simply responded according to the body posture, would have been removed from the analysis due to an inability to fit a PF to the data.

Linking the body-context effect to brain development, we found that the extent of influence of body context on facial expression perception was significantly related to the microstructure of fibre tracts between pSTS_body_ and ATL. Specifically, greater 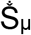 and R1 for this tract was associated with a smaller influence of body posture on facial expression perception. Importantly, this relationship was not significant after removing the variability associated with age, suggesting that the behaviour-structure association is mainly driven by factors relating to brain maturation, rather than age-independent characteristics influencing perception. While it is unclear how microstructural characteristics, indexed by 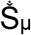 and R1, in tracts linking pSTS_body_ and ATL leads to less influence of body on facial expression perception, the location of these tracts lends support to previous findings in humans and monkeys pointing towards ATL as potential sites of face and body integration(Fisher & Freiwald, 2015; Harry et al., 2016; Teufel et al., 2019). Indeed, within the context of recent proposals put forward by Taubert et al (2021), our microstructural results provide some support for face and body networks being weakly integrated in early stages of processing, insofar as the link between tract microstructure from OFA to FFA and body posture perception indicates some processing of body posture within face pathways, with face and body processing becoming more integrated downstream of FFA, particularly leading to ATL.

More generally, our findings speak to the organisation of face processing in the visual system by providing support for revised face perception models (Duchaine & Yovel, 2015; Grill-Spector et al., 2018). While older models argued for separate pathways for facial identity and facial expression (Bruce & Young, 1986; Haxby et al., 2000) with FFA and STS thought to be specific to identity and expression processing, respectively, our results support more recent findings suggesting that FFA plays a role in the processing of facial expression (Bernstein & Yovel, 2015). Specifically, we show that the microstructure of tracts linking OFA to FFA are related to the development of facial expression discrimination ability, while none of the STS-related tracts show this association. It is worth highlighting that our analyses were restricted to core face and body processing regions, as we were particularly interested in the integration of face and body signals, with ATL as a likely site of possible integration. However, it is clear that brain areas beyond core face perception regions, such as the amygdala, play a critical role in emotion perception (Phelps & LeDoux, 2005). A deeper understanding of the relationship between facial expression perception and brain structure during development will therefore necessitate an exploration of the wider face and body perception networks in future.

The results of our study have to be considered within the context of a key limitation being that we focussed on only two emotional expressions: anger and disgust. The decision to focus on a limited number of emotions was partly based on feasibility constraints, dictated by the use of psychophysical methods. We specifically chose these two emotions for several reasons: first, disgust is one of the last emotions to be reliably recognised during development, maximising the likelihood of observing substantial developmental change across both our child and adolescent participants. Second, a key aim of our study was to focus on both emotional face and body signals, as well as their integration. Previous research has shown a more robust effect of body context with anger and disgust compared to other emotions (Aviezer et al, 2008), and we wanted to maximise the potential influence of body context in order to be sensitive to individual differences across development. Finally, the emotions of disgust and anger are associated with body postures that are clearly identifiable. By contrast, other postures like sadness and happiness are less easily conveyed through a simple posture (Lopez et al., 2017). While the limitation to these two emotions is important to keep in mind, there is no reason to believe that the key principles identified here, that facial expression precision drives the influence of body posture, as well as the relationships to microstructure, would not also apply to other emotions.

In summary, we focussed on the well-defined face and body networks to study the interplay between development of brain structure and perception in children and adolescents. We show that facial expression and body posture perception have a protracted developmental trajectory, extending far into adolescence. We find a similar developmental profile for the integration of face and body signals and demonstrate that the changing influence of body posture on facial expression can largely be explained by improvements in facial expression discrimination. We demonstrate that different microstructural characteristics of tracts within face and body pathways differentially relate to variability in facial expression discrimination in an age-dependent or age-independent manner. In particular, myelination and changes to intracellular signal-dominated microstructure, e.g. complexity of neuronal and glial processes, are predominantly associated with age-dependent improvements in facial expression and body posture perception. A more general, composite measure of microstructure, FA, is mainly associated with age-independent differences in perceptual performance. Overall, our results highlight the protracted development of facial expression perception, body posture perception, and context effects on facial expression perception. Moreover, our results shed light on the constraints that white matter microstructure imposes on behaviour and highlights the utility of using complementary measures of microstructure to study the links between brain structure, function, and perception.

## Acknowledgments

The authors would like to thank John Evans, Slawomir Kusmia, Allison Cooper and Sila Genc for their support with data acquisition. We would also like to thank the participants and their families for participating in the study.

